# Development of novel high-resolution melting markers for the assessment of hybridization between *Mauremys sinensis* and *Mauremys reevesii* in South Korea

**DOI:** 10.1101/2023.12.21.572907

**Authors:** Hae-jun Baek, Eujin Cheong, Youngha Kim, Kyo Soung Koo, Su-Hwan Kim, Chang-Deuk Park, Ju-Duk Yoon

## Abstract

The Chinese striped-necked turtle *Mauremys sinensis* was first reported in the wild in South Korea in 2012. Owing to its potential to hybridize with the native and endangered *Mauremys reevesii* in South Korea, leading to ecosystem disturbance, *M. sinensis* was designated as an ecosystem-disturbing species (EDS; hereafter) in 2020. Such hybridization between *M. sinensis* and *M. reevesii* has previously been reported in Taiwan. To protect and conserve the native South Korean *M. reevesii* population, the extent of its hybridization needed to be examined. Therefore, we developed novel high-resolution melting markers to elucidate the status of hybridization between *Mauremys sinensis* and *Mauremys reevesii* in South Korea. We identified the presence of *M. sinensis*, *M. reevesii*, and their hybrid across 47 sites in South Korea. Among these sites, five locations were confirmed as areas of sympatry for both *M. sinensis* and *M. reevesii*, with one hybrid individual identified at the Jeonju-si site. In total, hybrids were identified at four different sites. For more definitive species identification, blood samples were collected from the specimens and subjected to mitochondrial DNA (mtDNA) cytochrome c oxidase subunit I (*COI*) gene and nuclear DNA (nDNA) *R35* gene analysis, leading to the identification of two more hybrids from two other sites. To date, no hybrids of *M. sinensis* have been confirmed to reproduce in the wild, yet this requires further research for validation. The high-resolution melting (HRM) marker in the *R35* gene developed in this study yielded three distinct curves that were designated as *M. sinensis*, *M. reevesii*, and *M. sinensis* × *M. reevesii*, aligning with those of the *R35* gene sequencing. The novel markers developed in this study serve as a valuable tool that can be used in future research on rapid detection of hybridization.

## Introduction

The freshwater Chinese striped-necked turtle *Mauremys sinensis* (order Testudines; family: Geoemydidae) is listed as Endangered on the IUCN Red List and CITES Appendix III. This species is native to mainland China (Fujian, Guangdong, Guangxi, Hainan, and Zhejiang provinces), Taiwan, and Vietnam and has been introduced to South Korea, the United States (Florida), Europe, and Japan [1]. Although the specific dates and purposes of introduction remain unknown, it is believed that *M. sinensis* was imported as a substitute for *Trachemys* spp. (red-eared sliders), which were banned from import and distribution when designated as ecosystem-disturbing species in 2001 [2].

*Mauremys* is a globally traded group that was newly identified in the wild in South Korea in 2012 [3] (pp. 23-26), and its establishment in the country was confirmed in 2016 by Lee et al. [4]. Adult *M. sinensis* individuals are approximately 25 cm in length, with females growing larger than males [2]. They have a recorded maximum lifespan of 22.8 years in captivity and are omnivorous with distinct prey preferences observed between males and females. Adult females are omnivorous with a notable herbivorous preference, whereas adult males and young individuals are carnivorous [2].

Geoemydidae exhibit high levels of interspecific (intrageneric) hybridization, which has been reported to occur frequently in captivity owing to the high demand for this species as pets. When these human-created hybrid individuals are released into the wild, they pose a substantial a risk of introgression, potentially leading to the loss of unique genetic traits within wild populations. According to Stuart and Parham [5], 14 new turtle species have been reported in Geoemydidae in the last 20 years, six of which have been shown to be a result of hybridization in captivity. The hybridization of *M. sinensis* with other species of *Mauremys* has been reported, resulting in the creation of hybrids such as *M. sinensis* × *M. reevesii*, *M. sinensis* × *M. annamensis*, and *M. sinensis* × *C. trifasciata* [5–9].

The semi-aquatic turtle *Mauremys reevesii* is native to South Korea is protected by law as Natural Monument No. 453 and is listed as Endangered Wildlife II in South Korea [10]. Adults of the species can grow up to 30 cm in length [11]. This species can be distinguished from *M. sinensis* by its dark brown carapace with three ridges and lemon-yellow, c-shaped mark on the side of its face. Owing to habitat loss and destruction, the distribution of *M. reevesii* is limited. Recently, non-native turtle species such as *M. sinensis* have been introduced into its range, forcing them into coexistence [2,12–16].

Notably, habitat surveys are urgently needed in South Korea to proficiently determine the effect of invasive species on local fauna, which will allow for the development of more effective conservation strategies for native species. Therefore, in the present study, we aimed to identify the habitats of *M. sinensis* and *M. reevesii* and perform a clear molecular analysis for species identification. In addition, we developed a novel genotyping marker in the *R35* gene using high-resolution melting (HRM) analysis. HRM analysis facilitates genotyping by determining DNA sequence variations, such as single-nucleotide polymorphisms (SNPs), based on the melting temperature (Tm) of real-time PCR products [17–19]. This method is more user-friendly and cost-effective compared to standard sequencing methods and microsatellite marker analysis [20–22]. Such methods have been widely used in hybridization studies and for species identification [23–28]. Therefore, in addition to surveying *M. sinensis* and *M. reevesii,* we aimed to develop HRM markers that could distinguish these species and their hybrids for future genetic monitoring.

## Materials and methods

### Distribution of M. sinensis, M. reevesii, and their hybrid

A survey of the distribution of *M. sinensis* in South Korea and its sympatric sites with *M. reevesii* was conducted from 2017–2022. Survey sites were selected based on the National Institute of Ecology’s reports, namely the “Nationwide Survey of Non-native Species in Korea from 2016 to 2021 [13–16,29]” and the 2018 report, “Investigating Ecological Risk of Alien Species [2].” In total, 48 sites were selected, as listed in Table 1. Field surveys were conducted using binoculars (Swarovski, Absam, Tyrol, Austria, SLC 8×42 WBHD) and a field scope (Swarovski, ATM 65HD), and identification was based on external features such as three distinctive ridges on the carapace and side lines on the face. [6]. Most survey sites were located in urban centers and included ecological parks, reservoirs, and rivers close to municipalities, schools, and places of worship. Given the basking nature of turtles, surveys were conducted during the day.

**Table 1.**
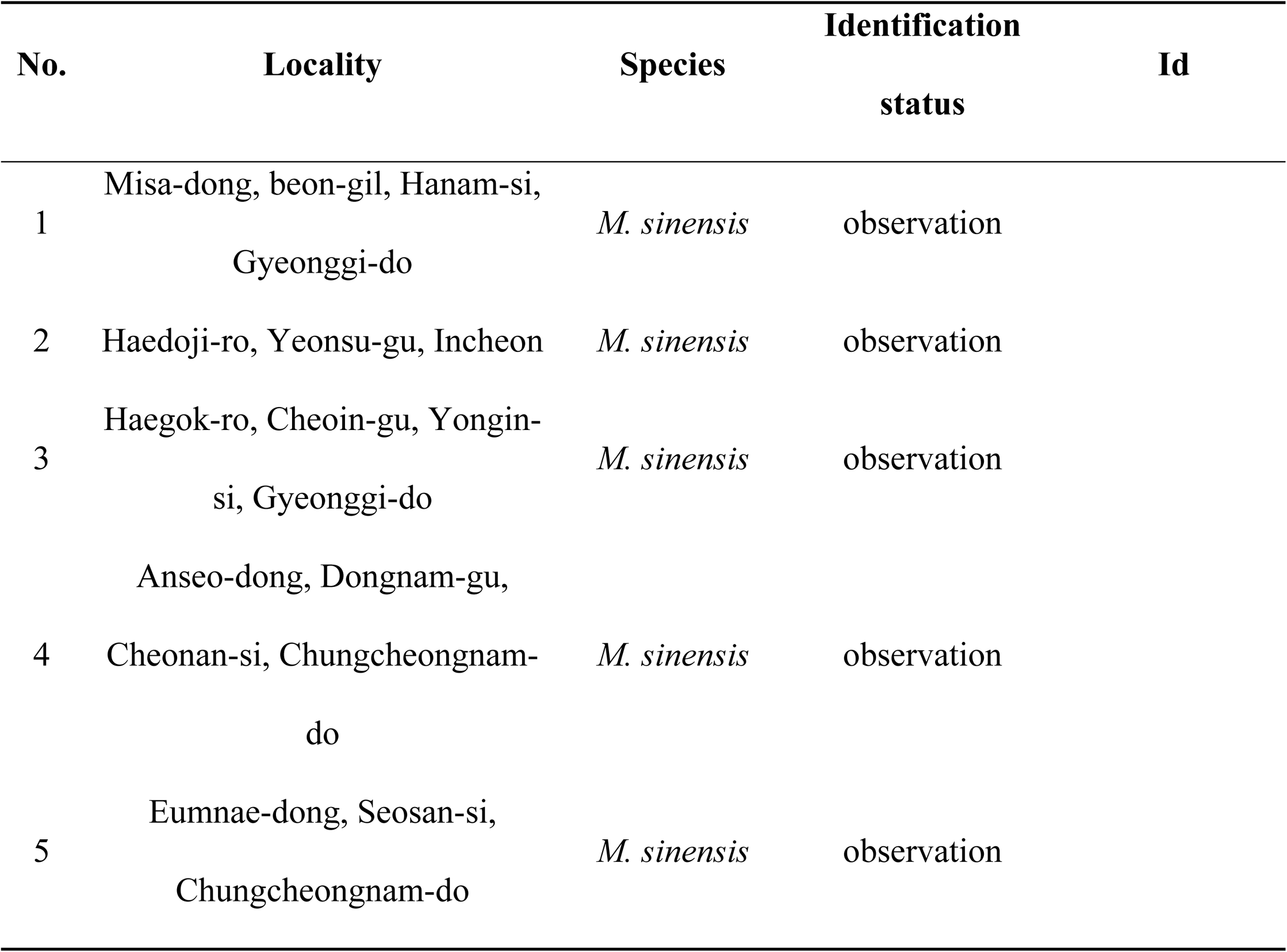
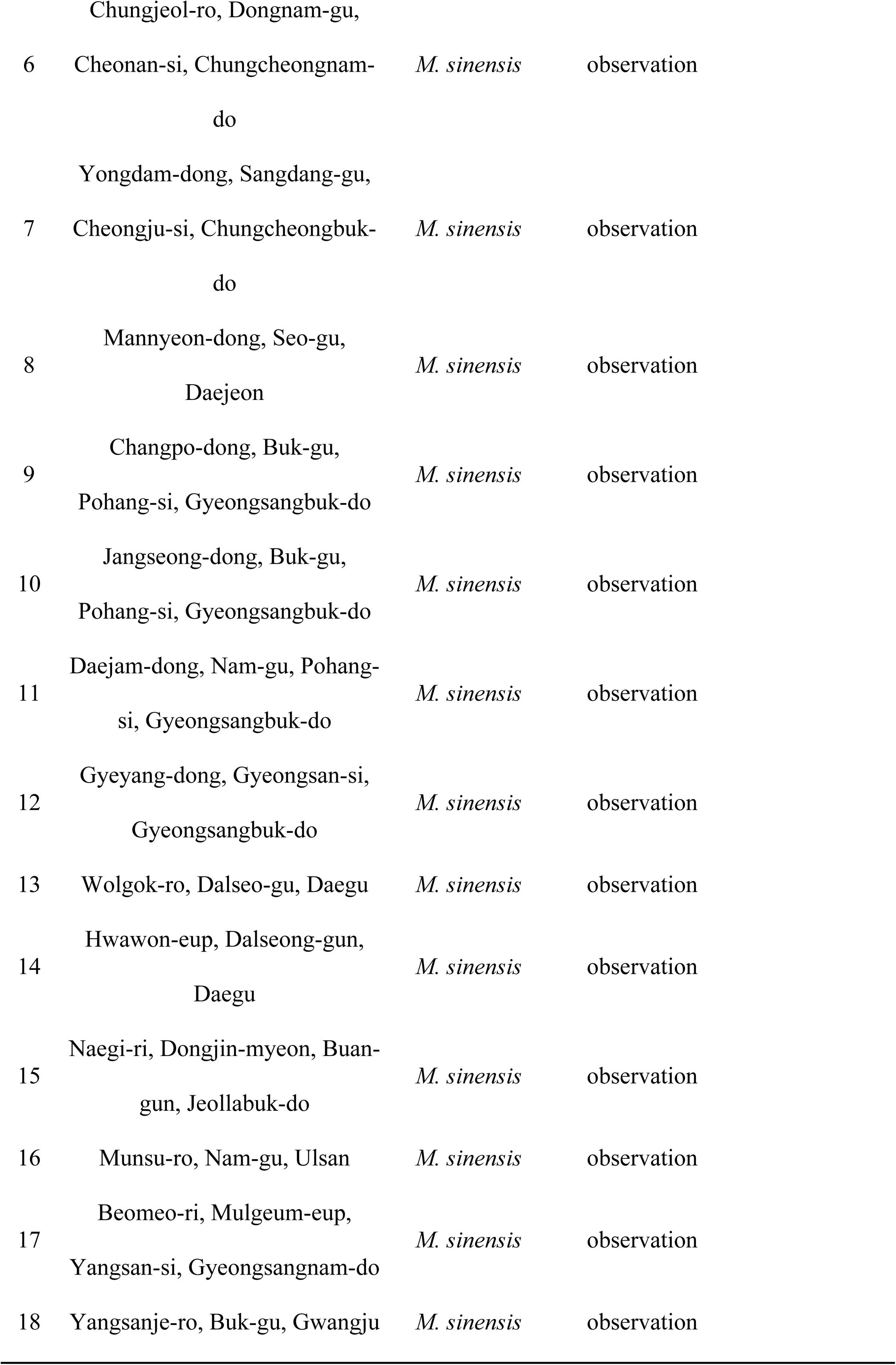
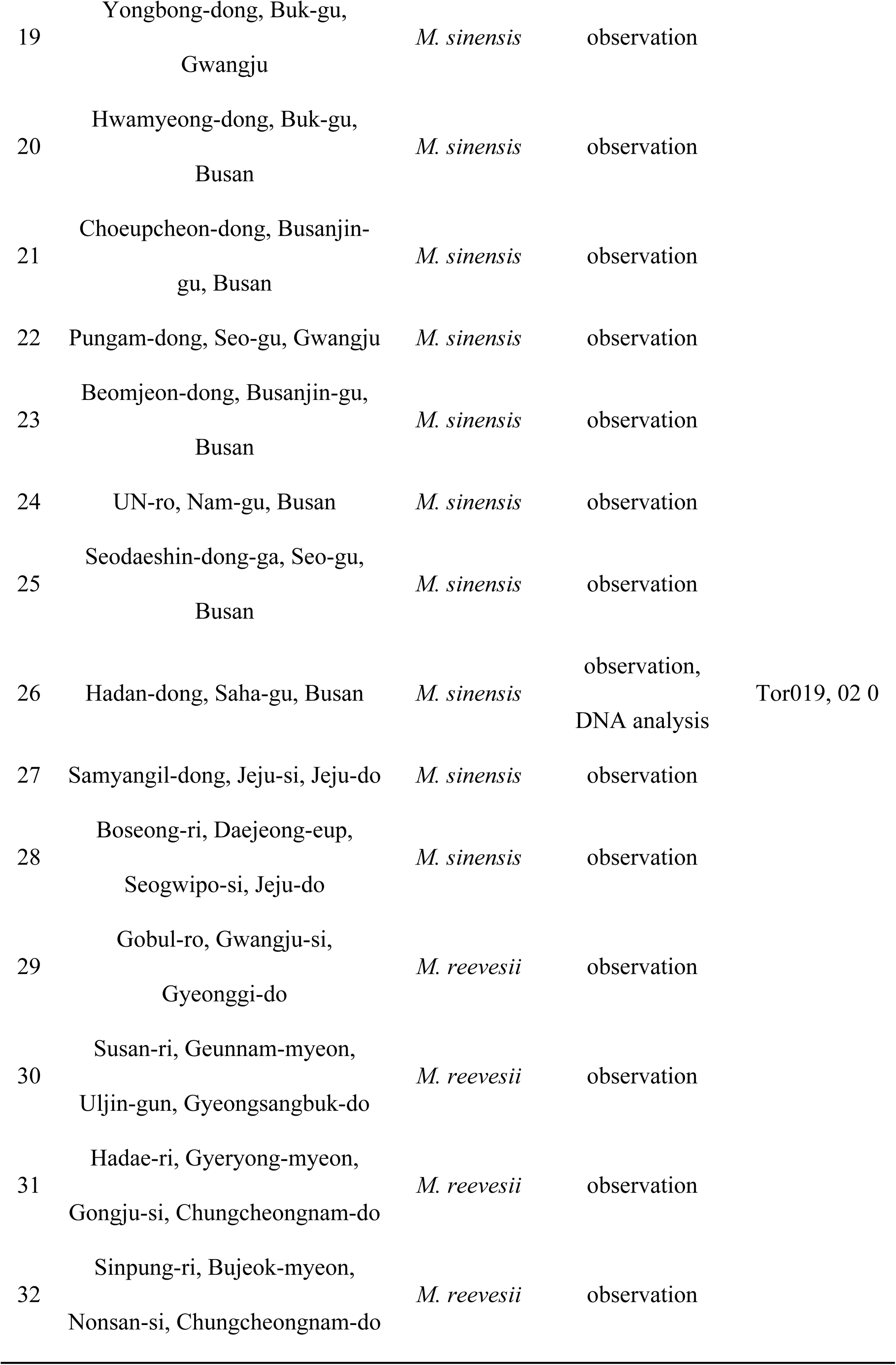
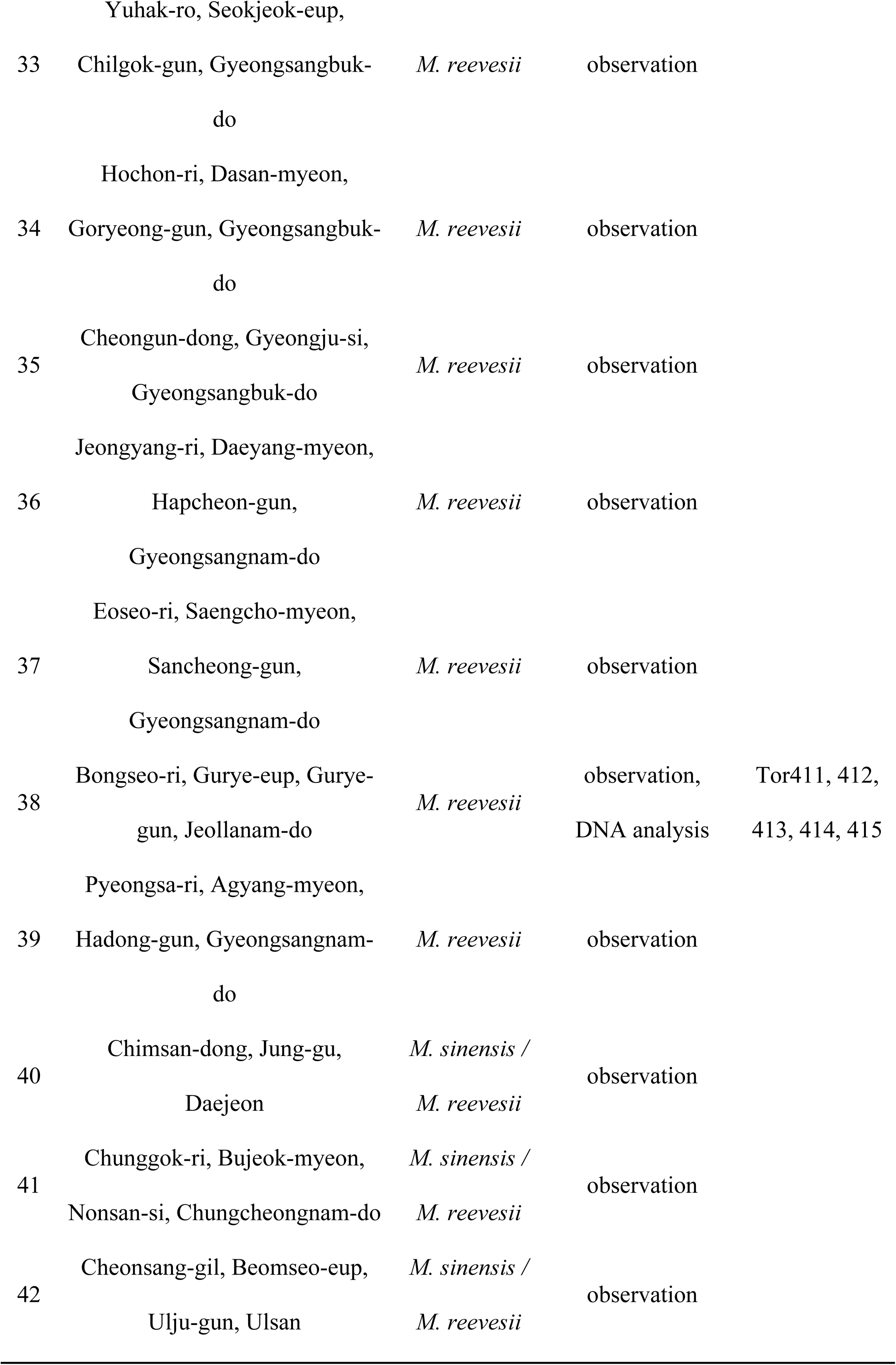
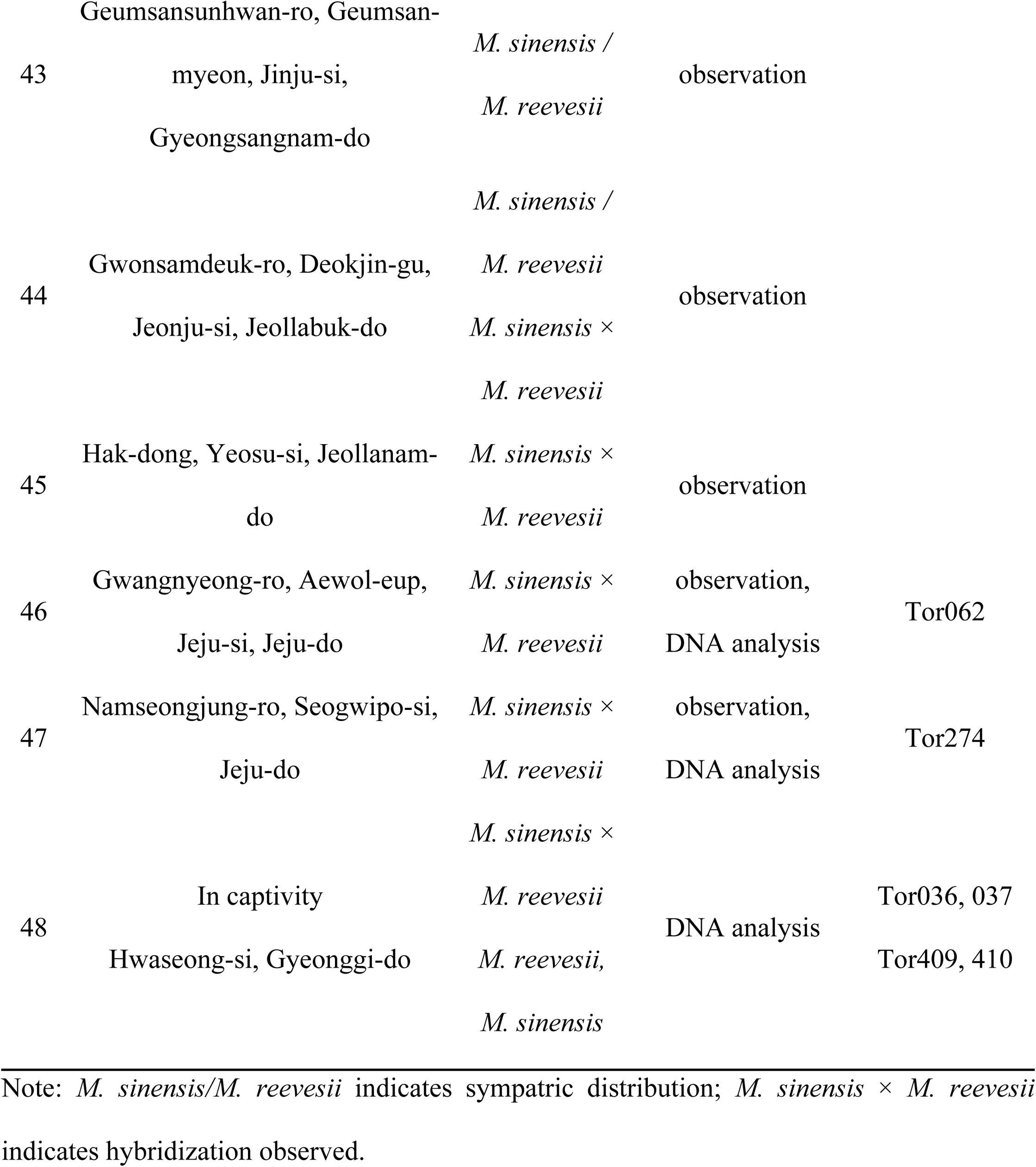
Survey sites of *Mauremys* species in South Korea, species found, identification methods, and individuals used for genetic analysis.

### Hybrid specimen sampling

To confirm hybridization between *M. sinensis* and *M. reevesii* in both wild and in captivity, we visited *M. reevesii* breeding farms in South Korea in 2018 (Hwaseong-si, Gyeonggi-do). Two suspected hybridized individuals were identified in this captive facility, and their blood samples were collected. To examine the presence of hybrids in the wild, we have been conducting capture surveys since 2020 using turtle traps at locations where habitat surveys have indicated that hybridized individuals may be present or where hybrid individuals have been recognized by the public (Jeonju-si, Jeollabuk-do, Yeosu-si, Jeollanam-do, Jeju-si, Jeju-do, Seogwipo-si, Jeju-do). Captured individuals were photographed, and their blood was drawn and re-radiated at the location of capture. The experiments were conducted following institutional guidelines and permission was obtained from the Animal Care and Use Committee of the National Institute of Ecology’s Center, South Korea (NIEIACUC-2021-020).

### Phylogenetic analysis

DNA was extracted using a Qiagen DNeasy blood and tissue kit (Qiagen, Hilden, Germany) following the manufacturer’s instructions. For molecular identification, the mitochondrial DNA cytochrome c oxidase subunit I (*COI*) gene and nuclear DNA *R35* genes were used. The novel primers used to amplify the target regions were designed in this study. The targeted gene of *COI* was amplified using MAU-MT-CO1-05190f 5′–TARTTAACAGCTAAAYACCC-3′ and MAU-MT-CO1-07039r 5′–AACCTATAATYTAACCTTGACAA-3′. For sequencing MAU-MT-CO1-05831f, 5′-TAACTATCTTTTCCCTYCACCTA-3′ was used. The *R35* gene of nuclear DNA intron 1 was amplified using MAU-R35-1f 5′-CAAAAGTCATTCTCTGGCTTC-3′, MAU-R35-2f 5′–GTCAGACTTCTTTGCATATTTGTAA-3′, and MAU-R35-1r 5′-CAACTATGTGCTGGACAG-3′ The AccuPower® PCR PreMix (BIONEER, Daejeon, South Korea) were used for both amplifications, and 20 ng of genomic DNA was obtained with a total volume of 20 µL. PCR conditions were as follows: 5 min at 95 ℃ for initial denaturation, 30 cycles of 20 s at 95 ℃ (denaturation), 20 s at 55 ℃ (annealing), 2 min at 72 ℃ (elongation), and final elongation for 5 min at 72 ℃. All samples were purified using an AccuPrep PCR Purification Kit (BIONEER) and sequenced on an ABI3730XL system (Applied Biosystems, Waltham, Massachusetts, USA) following the manufacturer’s instructions.

The sequences of both *COI* and *R35* genes were edited and aligned using Geneious 5.3.6 (BIOMATTERS, Auckland, New Zealand). Multiple sequence alignments were performed using CLUSTAL X [30]. Pairwise sequence divergence within and between different *Mauremys* species was estimated using MEGA version 7 [31]. The program Phase [32] was used to recover the alleles for the parental species.

DNA molecular phylogenies were reconstructed using the maximum likelihood method with default priors and 1,000 bootstrap replicates using MEGA version 7 [31]. The best-fit model for sequence evolution was selected using the same program. The best-fit model for *COI* based on the Bayesian information criterion was the T92 + G model, and that for *R35* was the T92 + G + I model. To ensure accurate reconstruction of the phylogenetic trees, 12 sequences of *M. sinensis*, 6 sequences of *M. reevesii*, 3 sequences of *Mauremys japonica*, and 5 sequences of *Mauremys mutica* were used, and *Cuora aurocapitata* (AY874540) and *C. amboinensis* (NC_014769) were used as the outgroups.

### Development of HRM markers

To design a primer combination for PCR amplification of the HRM molecular marker in the intron 1 region of the nuclear *R35* gene of *M. sinensis* and *M. reevesii*, we analyzed the DNA sequence data of species of the genus *Mauremys* from the NCBI GenBank database (DQ386678, JN860581, GQ259469, GQ259459, HQ442372, GU085687, JN860580, MK726303, DQ386656, MT961449, HQ442386, KX374280, MT961470, MT961473, and MT961487). These sequences were aligned with the newly deciphered DNA sequence data using ClustalW in BioEdit 7.2.5 [33], and a DNA sequence matrix was created. Based on these results, ProFlex PCR System (Thermo Fisher Scientific, Waltham, MA, USA) was used to design new PCR primer combinations containing SNPs that could distinguish *M. sinensis*, and *M. reevesii,* and their hybrids. The parameters included the length of the oligonucleotide sequence within 18–20-mer, the GC content between 40–55%, and the Tm value between 50–60 °C. The novel PCR primer sets for HRM analysis were designed as MAU-R36-0658f (5′–TCAGCTTCTCAGCTTCTTTC-3′) and MAU-R36-0713r (5′-CAGTGCCAGGCAGGATT-3′) (Fig 1). PCR amplification was carried out using the QuantStudio5 Real-Time PCR System (Thermo Fisher Scientific Waltham, Massachusetts, USA) with MeltDoctor HRM Master Mix (Thermo Fisher Scientific) to perform HRM analysis. The holding stage consisted of an enzyme-activation step at 95 °C for 10 min, and the cycling stage consisted of a 40-cycle denaturation step at 95 °C for 15 s and an annealing/elongation step at 60 °C for 1 min. The melt curve/dissociation stage consisted of denaturation at 95 °C for 10 s, annealing at 60 °C for 1 min, HRM at 95 °C for 15 s, and annealing at 60 °C for 15 s.

**Fig 1.**
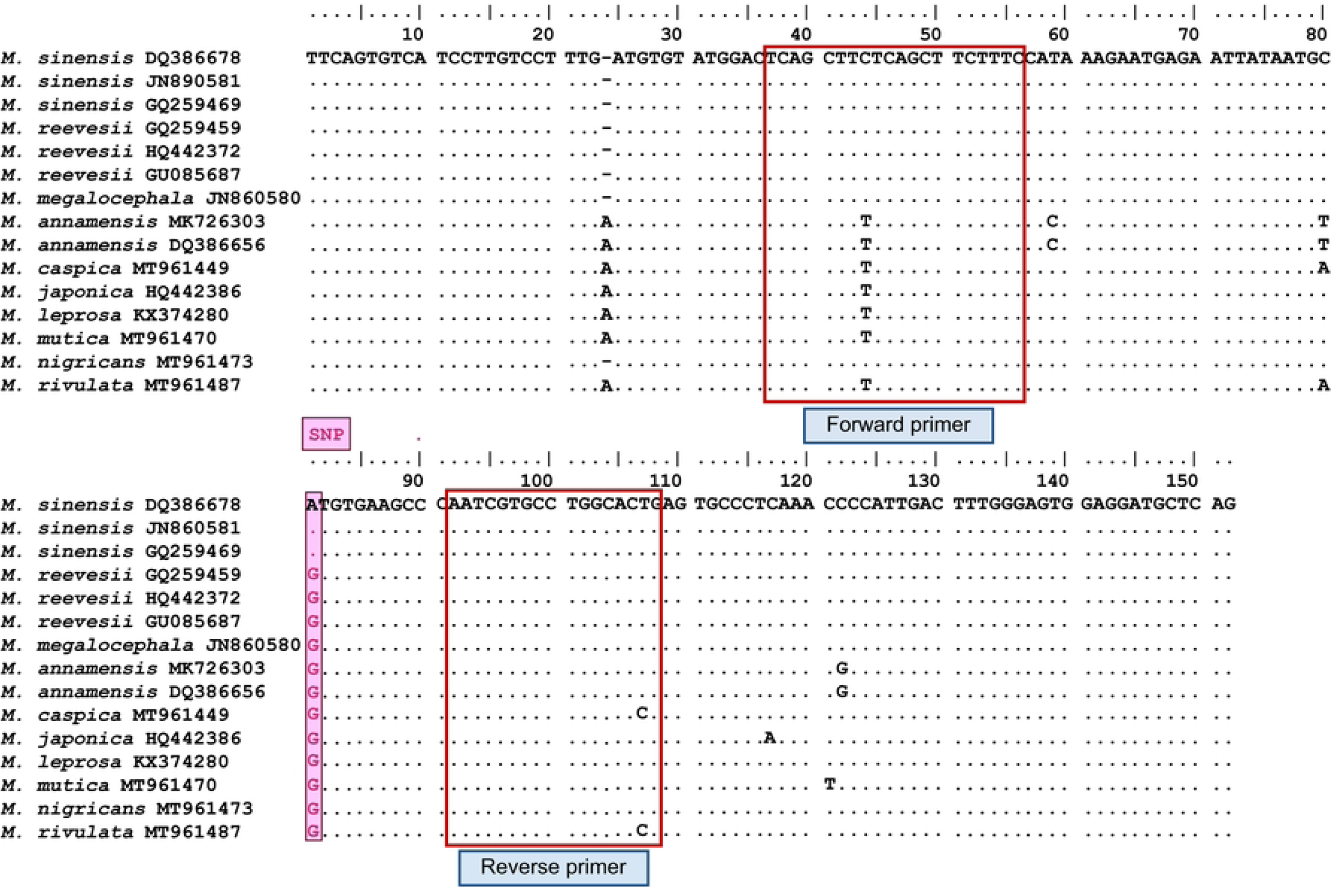
DNA sequence matrix of the intron I region of the nuclear *R35* gene containing single-nucleotide sequences (SNPs) and PCR primer combinations that can distinguish *Mauremys sinensis*, *M. reevesii,* and their hybrids.

## Results

### Distribution of M. sinensis, M. reevesii, and their hybrids

We confirmed the presence of *M. sinensis*, *M. reevesii,* and their hybrid at 47 sites across South Korea (Table 1, Fig 2). At five of these sites, *M. sinensis* and *M. reevesii* coexisted, with the identification of a hybrid individual occurring at one site. The sympatric sites for the two species were Daejeon (Chimsan-dong), Nonsan (Cheonggok-ri), Ulju (Cheonsang-gil), Jinju (Jangsari), and Jeonju (Kwon Samdeok-ro), with a hybrid individual identified in Jeonju. In total, the hybridization of *M. sinensis* with *M. reevesii* was confirmed at four sites: Jeonju (Kwon Samdeok-ro), Yeosu City (Hakdong), Jeju City (Gwangnyeong-ri), and Seogwipo City (Namjung-ro). *Mauremys sinensis* was confirmed at 33 sites across the country, whereas *M. reevesii* was restricted to 16 sites. *Mauremys sinensis* was mainly found in urban ecological parks with high human traffic, whereas *M. reevesii* was more likely to be found in less-disturbed natural habitats, such as rural areas.

**Fig 2.**
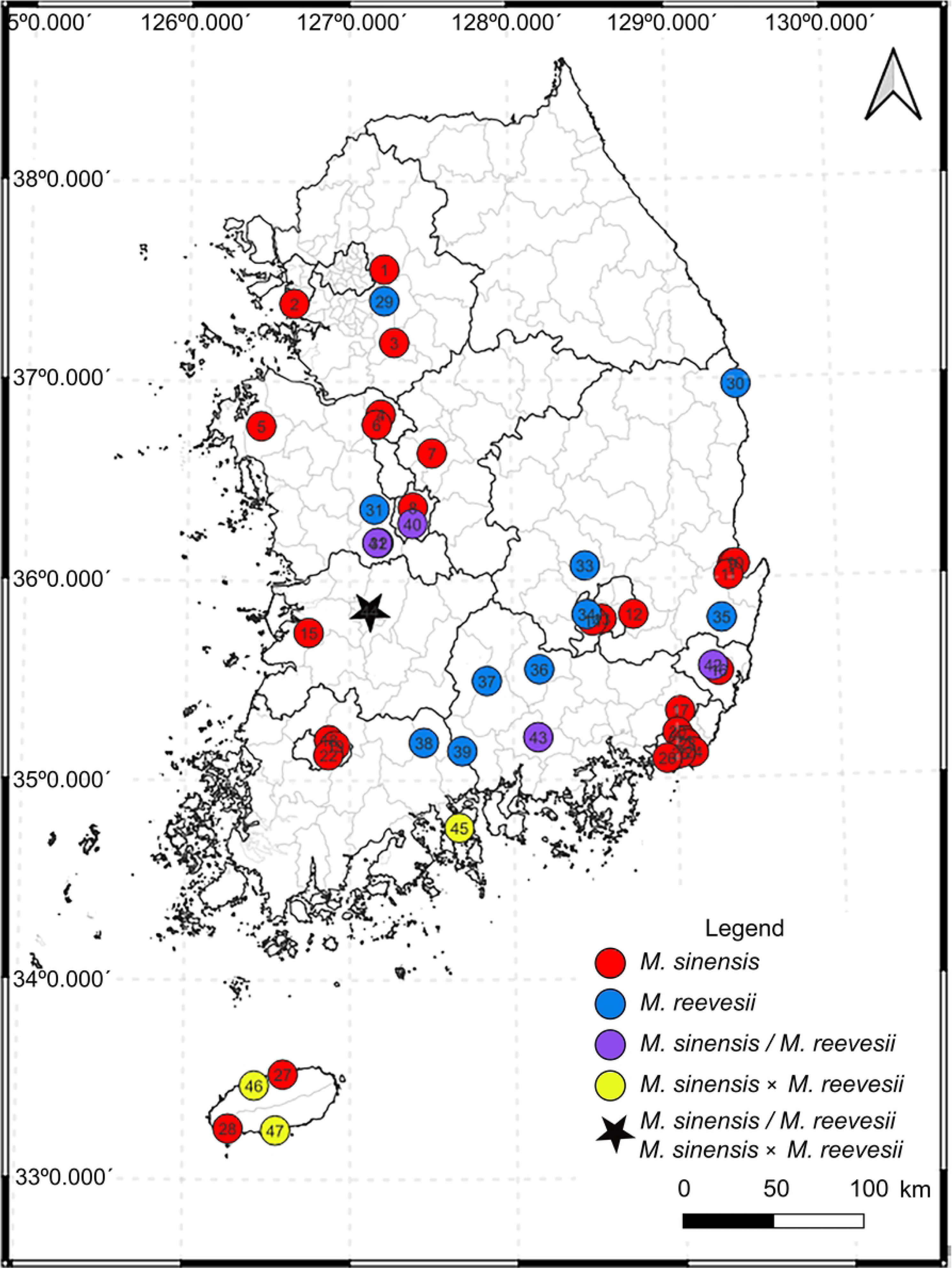
Survey sites of *Mauremys* species in South Korea and species information.

### Occurrence of M. sinensis × M. reevesii hybrids

During the field survey, traps were set to capture turtles at six of the seven sites where *M. sinensis* and *M. reevesii* coexisted or where hybrid individuals were visually identified, excluding Ulju-gun. The traps were made of transparent acrylic measuring 70 × 70 cm, with a depth of 20 cm, and the bait included was Japanese *sardinella* (fish) or pork. Nine turtles were captured, of which two were *M. sinensis*, five were *M. reevesii*, and two were hybrids (Fig 3).

**Fig 3.**
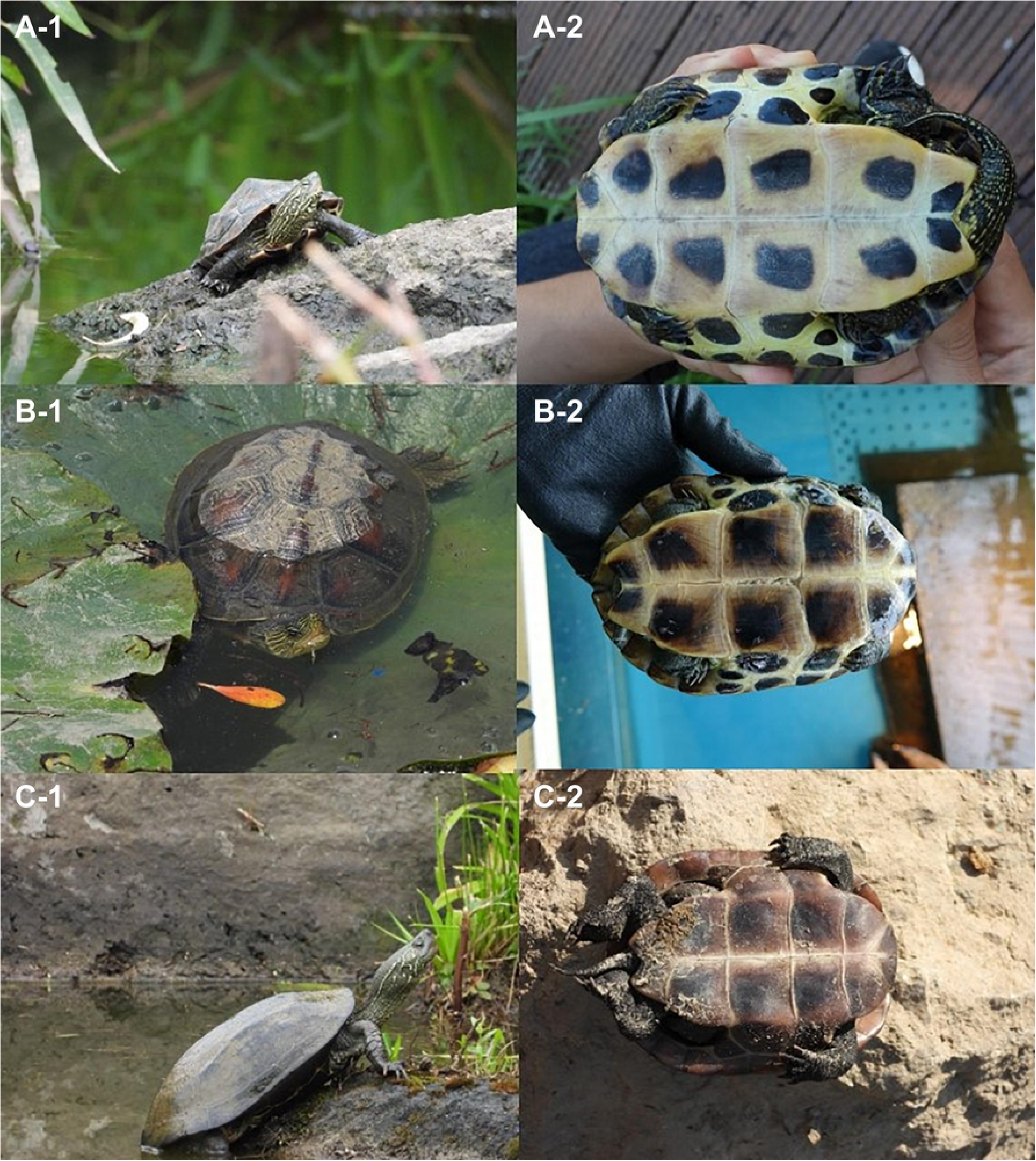
External differences among *Mauremys sinensis*, *M. reevesii*, and their hybrid; (A–1) Hybrid found on Jeju Island, (A-2) plastron of hybrid, (B-1) *M. sinensis*, (B-2) plastron of *M. sinensis*, (C-1) *M. reevesii,* and (C-2) plastron of *M. reevesii*.

### Phylogenetic analysis

The nine individuals captured in turtle traps and the four individuals captured from captivity were used for phylogenetic analysis. A total of 24 sequences (11 from GenBank, including two outgroups belonging to *Cuora*) were included for *COI* analysis and 31 sequences (18 from GenBank) for *R35* analysis. The partial sequence (615 bp) of the mtDNA *COI* gene from the 13 specimens represented two distinct clades in the maximum likelihood reconstruction (Fig 4). The pairwise sequence divergence between different clades ranged from 3.6% ± 0.7% to 4.4% ± 0.8%, and the overall sequence divergence was 2.6% ± 0.4%. The four suspected hybrids (Tor 036, 037, 062, and 274) belonged to the *M. sinensis* and *M. reevesii* clades. Tor 036 belonged to *M. sinensis* whereas as all other suspected hybrids belonged to *M. reevesii* (Table 2). The *R35* gene (863 bp) from 31 sequences comprised three separate clades: *M. sinensis*, *M. reevesii*, and *M. japonica* (Fig 5). The overall sequence divergence was 0.5% ± 0.1%, and that between *Mauremys* species ranged from 0.4% ± 0.1% to 1.1% ± 0.3%. All suspected hybrid individuals possessed both *M. sinensis* and *M. reevesii* alleles, indicating that they had formed through independent hybridization events.

**Fig 4.**
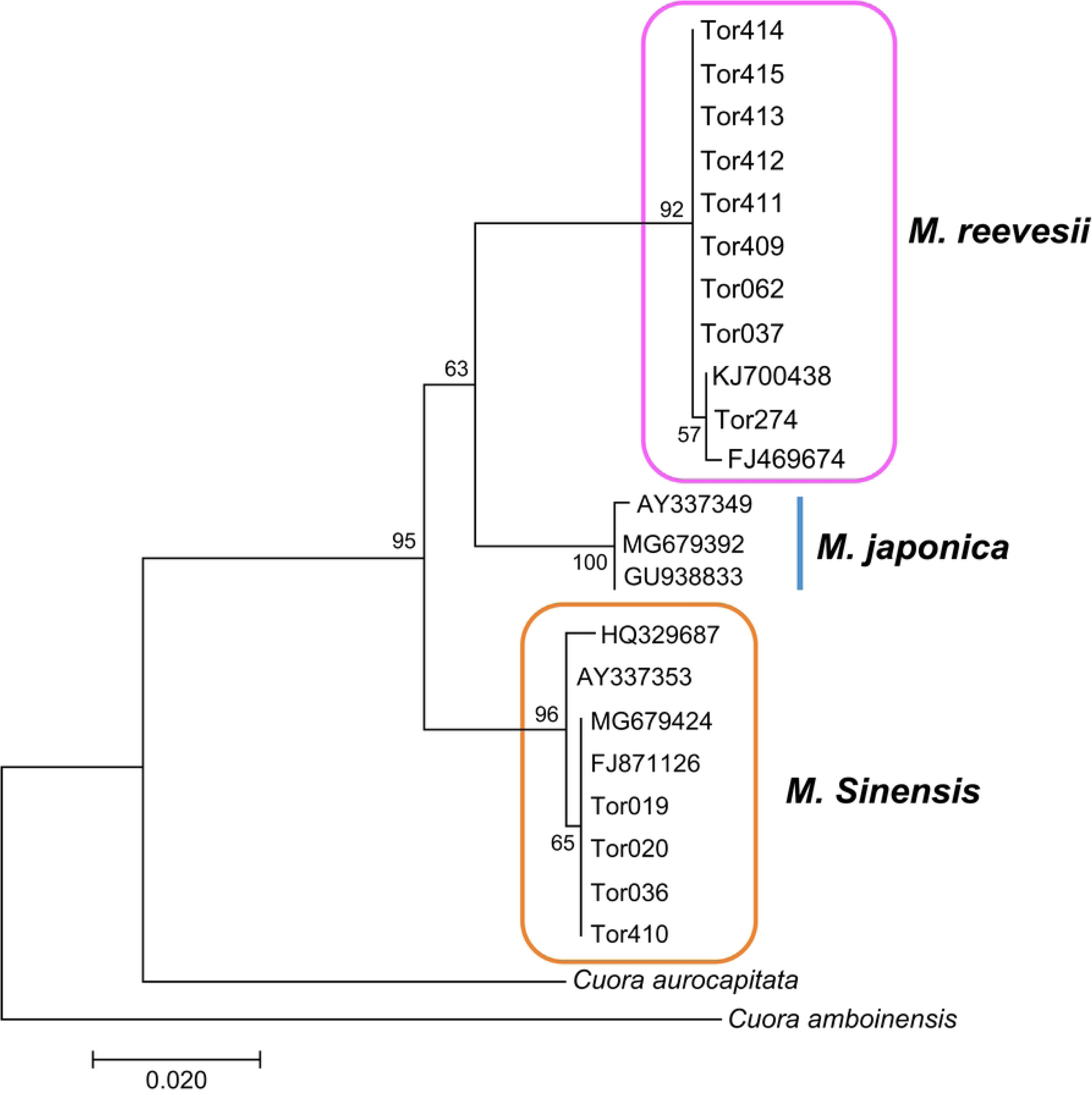
Maximum-likelihood tree based on an analysis of 22 *Mauremys* individuals using 615 bp of mitochondrial *CO* I sequences, including two *Cuora* species (*C. aurocapitata* AY874540, *C. amboinensis* NC_014769), with T92 + G and bootstrapping (1000 replications).

**Fig 5.**
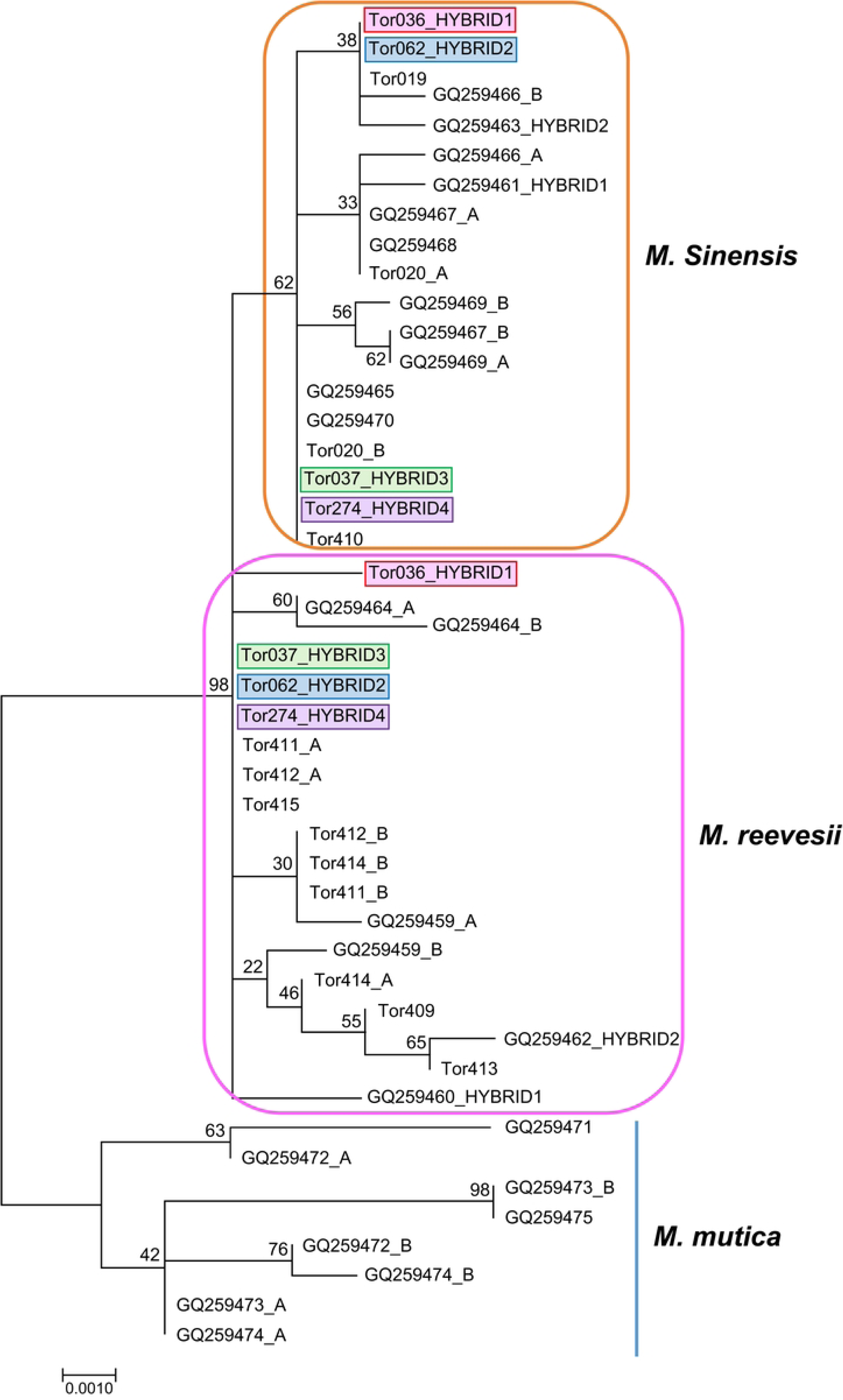
Maximum-likelihood tree based on an analysis of 31 *Mauremys* individuals using 863 bp of nuclear *R35* sequences, with T92 + I + G and bootstrapping (1000 replications).

**Table 2.** Information on species, results of *CO1* and *R35* phylogenetic analysis, SNP sites of the *R35* gene based on the nucleotide sequence analysis, melting temperature (Tm), and species identification based on the high-resolution melting (HRM) analysis.

| No. | ID | Species | *COI* tree | *R35* tree | SNP | T<sub>m</sub><br>(°C) | HRM analysis |
| --- | --- | --- | --- | --- | --- | --- | --- |
| 1 | Tor019 | *M. sinensis* | *M.*<br>*sinensis* | *M. sinensis* | A | 74.9 | *M. sinensis* |
| 2 | Tor020 | *M. sinensis* | *M.*<br>*sinensis* | *M. sinensis* | A | 74.9 | *M. sinensis* |
| 3 | Tor036 | *M. sinensis* ×<br>*M. reevesii* | *M.*<br>*sinensis* | *M. sinensis* ×<br>*M. reevesii* | R | 75.2 | *M. sinensis* ×<br>*M. reevesii* |
| 4 | Tor037 | *M. sinensis* ×<br>*M. reevesii* | *M.*<br>*reevesii* | *M. sinensis* ×<br>*M. reevesii* | R | 75.2 | *M. sinensis* ×<br>*M. reevesii* |
| 5 | Tor062 | *M. sinensis* ×<br>*M. reevesii* | *M.*<br>*reevesii* | *M. sinensis* ×<br>*M. reevesii* | R | 75.2 | *M. sinensis* ×<br>*M. reevesii* |
| 6 | Tor274 | *M. sinensis* ×<br>*M. reevesii* | *M.*<br>*reevesii* | *M. sinensis* ×<br>*M. reevesii* | R | 75.2 | *M. sinensis* ×<br>*M. reevesii* |
| 7 | Tor409 | *M. reevesii* | *M.* | *M. reevesii* | G | 75.5 | *M. reevesii* |
|  |  |  | *reevesii* |  |  |  |  |
| 8 | Tor410 | *M. sinensis* | *M.* | *M. sinensis* | A | 74.9 | *M. sinensis* |
|  |  |  | *sinensis* |  |  |  |  |
| 9 | Tor411 | *M. reevesii* | *M.* | *M. reevesii* | G | 75.4 | *M. reevesii* |
|  |  |  | *reevesii* |  |  |  |  |
| 10 | Tor412 | *M. reevesii* | *M.* | *M. reevesii* | G | 75.7 | *M. reevesii* |
|  |  |  | *reevesii* |  |  |  |  |
| 11 | Tor413 | *M. reevesii* | *M.* | *M. reevesii* | G | 75.7 | *M. reevesii* |
|  |  |  | *reevesii* |  |  |  |  |
| 12 | Tor414 | *M. reevesii* | *M.* | *M. reevesii* | G | 75.6 | *M. reevesii* |
|  |  |  | *reevesii* |  |  |  |  |
| 13 | Tor415 | *M. reevesii* | *M.* | *M. reevesii* | G | 75.6 | *M. reevesii* |
|  |  |  | *reevesii* |  |  |  |  |

### Development of HRM markers

SNPs that could distinguish *M. sinensis*, *M. reevesii,* and their hybrid were examined in intron 1 of the nuclear *R35* gene, and novel molecular markers, including novel PCR primer combinations, were designed. The forward and reverse primers were highly conserved, with no genetic variation between *M. sinensis* and *M. reevesii*, and the SNPs showed a 1 bp genetic variation between the two species (Fig 1). Thirteen specimens were subjected to HRM analysis, all of which were divided into three genotypes: *M. sinensis* (A), *M. reevesii* (G), and *M. sinensis* × *M. reevesii* (R). The HRM analysis using the genotyping markers revealed that three specimens produced a single peak with Tm at 74.9 °C that corresponded to *M. sinensis* (A), six specimens with Tm at 75.4–75.7 °C corresponded to *M. reevesii* (G), and four specimens with Tm at 75.2 °C corresponded to *M. sinensis × M. reevesii* (G) (Table 2). Further analysis using the melting curves produced visibly different plot shapes (Fig 6). These results were consistent with the phylogeny of *R35* (Table 2).

**Fig 6.**
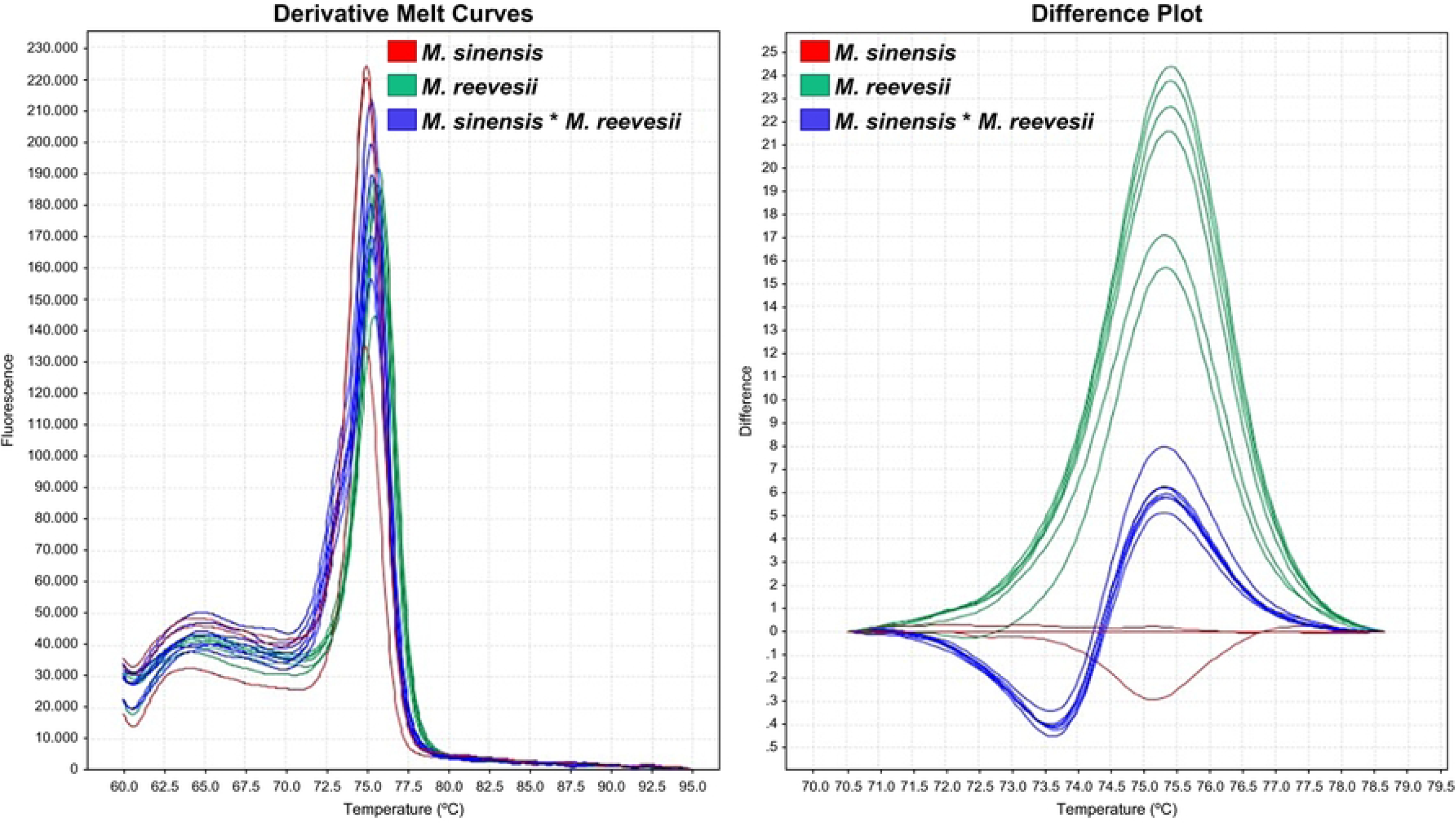
Derivative melt curves (A) and difference plot curves (B) produced by HRM analysis of *M. sinensis, M. reevesii*, and *M. sinensis × M. reevesii*.

## Discussion

In this study, we verified the occurrence locations of *M. sinensis* in South Korea, its sympatric areas with *M. reevesii*, and the first record of hybridization between *M. sinensis* and *M. reevesii* in the wild. The hybrid status of two suspected hybrids identified on Jeju Island were confirmed through genetic analysis, which revealed that *M. reevesii* was the maternal lineage and *M. sinensis* was the paternal lineage. Jeju Island has no reported native turtle species other than *Pelodiscus maackii*; however, five non-native turtle species have been recorded: *Trachemys scripta, Pseudemys concinna, Pelodiscus nelsoni*, *Mauremys sinensis,* and *Podocnemis unifilis* [29, 34]. The habitats where the hybrid individuals were recorded were unavailable for *M. sinensis* and *M. reevesii*, suggesting that the turtles identified in the present study were captive-bred individuals that were introduced into the wild rather than naturally reproducing individuals. However, further investigation is needed to confirm this.

In the present study, *M. reevesii* was identified in 16 sites, and 5 of the sites were areas of sympatry (Daejeon, Nonsan, Ulju, Jinju, and Jeonju) that contained both *M. reevesii* and *M. sinensis*. Based on our survey, we recommend that Deokjin Lake, located in Deokjin-gu, Jeonju-si, should be designated as an intensive management area because it is a registered area of sympatry of *M. sinensis* and *M. reevesii*. As hybrid individuals have been identified, the need for the active removal of turtles characterized as EDS (Ecosystem Disturbing Species; IAS in South Korea) is crucial. To date, the only non-native turtle species that have been documented to breed in the wild are *Trachemys* spp., *Pseudemys* spp., and *Chelydra serpentina*, with no reports of *M. sinensis* breeding. However, the breeding status of *M. sinensis* in South Korea is yet to be examined, thereby facilitating its urgent need.

Studies on the potential impacts of non-native turtles on native turtle populations include those by Jo et al. [35], Seo et al. [36], and Park [37] who confirmed that they have overlapping ecological statuses in terms of food sources and behavioral ranges. Non-native species disrupt native species diversity through competition, disease transmission, and hybridization with native species, increasing the risk of native species extinction and causing changes in ecosystem functions and service delivery [38].

Over the last 20 years, 161 tons of turtles have been imported into South Korea from 63 countries [39]. China, the country of origin of *M. sinensis*, is the largest domestic importer of turtles, with approximately 110 tons imported in the last two decades. The turtle business in China is extensive, with as many as 10 turtle farms in Hainan alone and different species of turtles reared in the same pond [40]. Indeed, *M. iversoni* (invalid) has also been identified as a hybrid arising from the co-rearing of *M. mutica* and *C. trifasciata* on turtle farms in China [40]. In particular, genetic pollution has been reported in *Mauremys*, with hybrids being frequently identified [6,8,41–45]. Ueno et al. [41] revealed a cross-back hybrid between *M. japonica* and *M. reevesii* and reported that the fertility and hatchability of F1s of *M. japonica* and *M. reevesii* were not significantly different from those of the parental species, posing a potential threat to the conservation status of *M. japonica*, which is endemic to mainland Japan. In addition, Lee et al. [9] reported that imported *M. sinensis* caused genetic introgression in native *M. reevesii* populations in the protected area, raising a critical risk to their conservation.

Previous studies on the management of non-native species in South Korea include Baek et al. [39], Kim and Koo. [46], Kim et al. [47], and Kim. [48]. These studies were primarily concerned with the policies and legislation that encompass both EDS and invasive species. However, improving laws and policies requires time and money, and practical management measures are needed immediately.

We propose to designate priority protected areas and removal points for EDS. The five sites identified in this study, which are crucial habitats for *M. reevesii*, should be designated as priority management areas, and seasonal management should be conducted considering the characteristics of the species. From May to July, when emergence and reproduction are concentrated, direct removal of EDS turtles using turtle traps and destruction of their nesting sites should be carried out. For *M. reevesii*, it is essential to estimate genetic pollution through genetic analyses and implement appropriate management measures. The HRM markers developed in the present study will be valuable for faster and cheaper genotyping of male and hybrid individuals. However, further research is required to develop eDNA markers specific to *Mauremys* for comprehensive nationwide habitat monitoring.

## Conclusions

This study was conducted at 48 sites across South Korea to identify hybrids of *M. sinensis* and *M. reevesii*. Hybrids were confirmed at four sites, with two individuals captured from Jeju Island. For further confirmation, genetic analysis was conducted, which verified that the maternal lineage of both individuals as *M. reevesii* and the paternal lineage as *M. sinensis*. In addition, two captive-bred hybrids were also recovered, suggesting that the hybrids identified in South Korea do not occur naturally, but are believed to be captive-bred individuals that were released. Among the sites identified in this study, Deokjin Park in Jeonju was designated as a priority management area for active EDS turtle removal activities, necessitating continuous monitoring and breeding studies in the future. We developed HRM markers capable of distinguishing between the two species and their hybrid, and were able to successfully distinguish between them. We expect that these markers will assist researchers in distinguishing between hybrids more rapidly and accurately.

## Acknowledgments

We would like to thank the members of the Invasive Alien Species Team at the National Institute of Ecology and the students at the Ewha Woman’s University for helping us to carry out the research.

